# Antisense, but not sense, repeat expanded RNAs activate PKR/eIF2α-dependent integrated stress response in C9orf72 FTD/ALS

**DOI:** 10.1101/2022.06.06.495030

**Authors:** Janani Parameswaran, Nancy Zhang, Kedamawit Tilahun, Devesh C. Pant, Ganesh Chilukuri, Seneshaw Asress, Anwesha Banerjee, Emma Davis, Samantha L. Schwartz, Graeme L. Conn, Gary J. Bassell, Jie Jiang

## Abstract

GGGGCC (G_4_C_2_) hexanucleotide repeat expansion in the C9orf72 gene is the most common genetic cause of frontotemporal dementia (FTD) and amyotrophic lateral sclerosis (ALS). The repeat is bidirectionally transcribed and confers gain of toxicity. However, the underlying toxic species is debated, and it is not clear whether antisense CCCCGG (C_4_G_2_) repeat expanded RNAs contribute to disease pathogenesis. Our study shows that C9orf72 (C_4_G_2_) antisense repeat expanded RNAs trigger the activation of the PKR/eIF2α-dependent integrated stress response independent of dipeptide repeat proteins that are produced through repeat-associated non-AUG initiated translation, leading to global translation inhibition and stress granule formation. Increased phosphorylation of PKR/eIF2α is also observed in the frontal cortex of C9orf72 FTD/ALS patients. Finally, only antisense (C_4_G_2_), but not sense (G_4_C_2_), repeat expanded RNAs can activate the PKR/eIF2α pathway. These results provide a mechanism by which antisense repeat expanded RNAs elicit neuronal toxicity in FTD/ALS caused by C9orf72 repeat expansions.

## Introduction

In 2011, GGGGCC (G_4_C_2_) hexanucleotide repeat expansion in the first intron of chromosome 9 open reading frame 72 (C9orf72) gene was identified as the most common genetic cause of frontotemporal dementia (FTD) and amyotrophic lateral sclerosis (ALS), two neurodegenerative diseases that are now believed to belong to a continuous disease spectrum with clinical, pathological, and genetic overlaps [1, 2]. In normal populations, the G_4_C_2_ repeat size is between two and thirty, whereas it expands to hundreds or thousands in FTD/ALS patients (referred to hereafter as C9FTD/ALS). C9FTD/ALS thus joins an increasing number of repeat expansion disorders including Huntington’s disease, myotonic dystrophy, and several spinocerebellar ataxias [3]. Based on initial pathological assessment of C9FTD/ALS patient postmortem tissues and lessons learned from other repeat expansion disorders, several pathogenic mechanisms by which expanded C9orf72 repeats can exert toxicity were proposed [4]. First, expanded G_4_C_2_ repeats inhibit C9orf72 mRNA transcription, leading to haploinsufficiency of C9orf72 protein [5, 6]; second, C9orf72 repeats are bidirectionally transcribed into sense G_4_C_2_ and antisense CCCCGG (C_4_G_2_) RNAs. These repeat-expanded RNAs may cause gain of toxicity by sequestering key RNA binding proteins into RNA foci and/or by production of toxic dipeptide repeat (DPR) proteins via non-canonical repeat-associated non-AUG-dependent (RAN) translation from all reading frames. More specifically, translating from sense G_4_C_2_ RNAs produces GA (Glycine-Alanine), GP (Glycine-Proline), and GR (Glycine-Arginine) DPR proteins, and translating from antisense C_4_G_2_ RNAs produces GP (Glycine-Proline), PA (Proline-Alanine), and PR (Proline-Arginine) DPR proteins [7]. In addition to these pure dimeric DPR proteins, there is also evidence of chimeric DPR proteins both in vitro and in patients [8-10].

How C9orf72 repeat expansions cause FTD/ALS has been extensively explored. Although reducing C9orf72 in zebrafish or *C. elegans* can cause motor deficits [11, 12], reduced or even complete deletion of C9orf72 in mice does not lead to FTD/ALS-like abnormalities, suggesting that loss of C9orf72 is not a main disease driver [13-19]. Supporting this, no missense or truncation mutations in C9orf72 are found in FTD/ALS patients yet [20]. On the other hand, several lines of studies, by expressing either G_4_C_2_ repeats [21, 22] or individual codon-optimized, ATG-driven DPR proteins [23-25], support that gain of toxicity from repeat expanded RNAs plays a central role in disease pathogenesis. Finally, loss of C9orf72, which plays a role in autophagy/lysosomal functions, can exacerbate toxicity from the repeat expanded RNAs [26, 27].

The underlying toxic species arising from C9orf72 repeat expanded RNAs that drive disease is still debated. Several RNA binding proteins (RBPs) are suggested to interact with G_4_C_2_ or G_4_C_2_ repeat RNAs and co-localize with RNA foci [28-35]. However, strong evidence supporting that loss of any proposed RBPs drives C9FTD/ALS is lacking. In contrast, ectopic expression of individual DPR proteins, especially GR and PR, causes toxicity in various model systems [23-25, 36-50]. To determine the relative contributions of RNA foci- and DPR protein-mediated toxicity, two studies employed interrupted repeats with stop codons in all reading frames to prevent DPR protein production and concluded that both sense and antisense RNAs are not toxic in *Drosophila* [51, 52]. This was challenged by another study showing both sense and antisense RNAs can cause motor axonopathy in zebrafish independent of DPR proteins [11]. Irrespective of RNA foci and DPR proteins, studies using antisense oligonucleotide (ASOs) to selectively degrade sense G_4_C_2_ repeat expanded RNAs strongly support its role in C9FTD/ALS pathogenesis. These sense strand-specific ASOs not only mitigate toxicity from C9orf72 repeat expansions in both transgenic mice expressing G_4_C_2_ repeats [22] and IPSC-derived neurons [28], but also reverse downstream cellular and molecular alterations such as nucleocytoplasmic transport deficits [53]. However, whether antisense C_4_G_2_ expanded RNAs contribute to C9FTD/ALS and thus are targets of intervention is less clear. Although PR translated from antisense strand is extremely toxic in model systems, PR or its aggregates are rare. Antisense RNA transcripts are also hard to detect in patient postmortem tissues. Surprisingly, several studies showed antisense RNA foci are as abundant as sense RNA foci in multiple brain regions [54-56], raising a possibility that antisense C_4_G_2_ repeat expanded RNAs also contribute to diseases [57]. In this study, we show that antisense C9orf72 C_4_G_2_ expanded repeats are neurotoxic independent of RAN translated DPR proteins. Antisense C_4_G_2_, but not sense, repeat expanded RNAs activate PKR/eIF2α-dependent integrated stress response, leading to global protein synthesis and stress granules formation. Moreover, the phosphorylation of PKR/eIF2α is significantly increased in C9FTD/ALS patients, suggesting that antisense C_4_G_2_ repeat expanded RNAs contribute to disease pathogenesis.

## Results

### C9orf72 antisense C_4_G_2_ expanded repeats are neurotoxic

To determine the contribution of C9orf72 antisense repeat expanded RNAs in FTD/ALS pathogenesis, we first generated a construct containing 75 C_4_G_2_ repeats using recursive directional ligation as previously described [24]. We included 6 stop codons (2 every frame) at the N-terminus to prevent unwarranted translation initiation and 3 protein tags in frame with individual DPR proteins at the C-terminus (**Fig. 1A**). Recent studies have shown that nucleotide sequences at 5’- and 3’-regions of expanded repeats regulate toxicity [58, 59]. Although the molecular mechanism of C9orf72 antisense transcription initiation is unknown, it has been shown that transcription can start from at least 450bp nucleotides upstream [60]. We therefore added 450bp of human sequence at the 5’-region of the antisense C_4_G_2_ repeats and termed this construct as “in_(C_4_G_2_)75”. When expressed in HEK293T cells, we detected abundant accumulation of antisense RNA foci, but not in control cells expressing 2 C_4_G_2_ repeats. Using antibodies against individual DPR proteins RAN translated from C_4_G_2_ expanded repeat RNAs or the protein tags in frame, we also observed production of GP, PR, and PA DPR proteins only in cells expressing in_(C_4_G_2_)75 but not 2 repeats (**Fig. S1A-B**). Antisense RNA foci and DPR proteins were also observed in mouse primary cortical neurons expressing in_(C_4_G_2_)75, but not in neurons expressing 2 repeats (**Figs. 1B and S1C**). Thus, in_(C_4_G_2_)75 produces antisense RNA foci and DPR proteins, two cellular pathological hallmarks observed in C9FTD/ALS patients.

**Figure 1.**
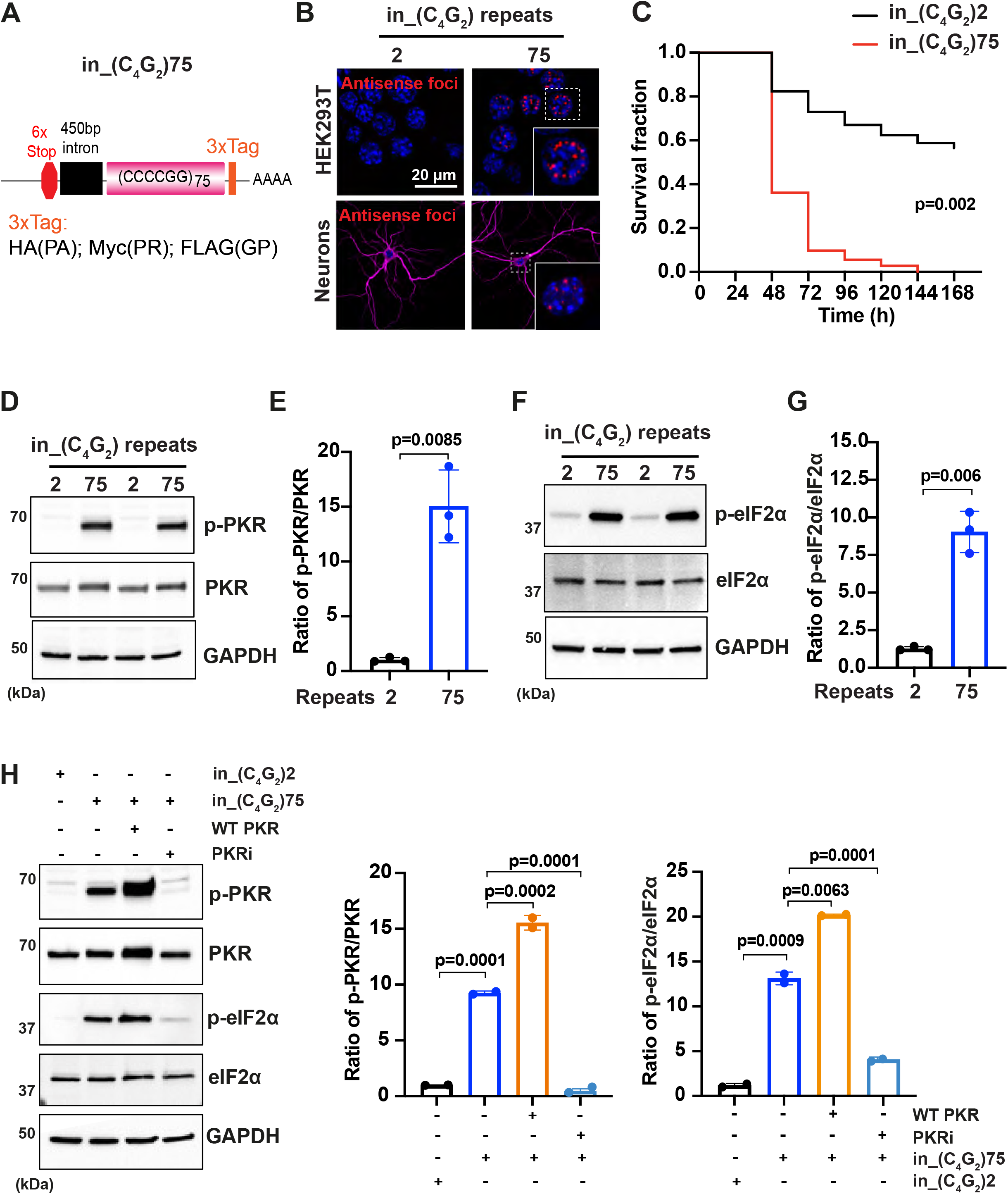
C9orf72 C_4_G_2_expanded repeats activate PKR/eIF2α-dependent integrated stress response and cause neuronal toxicity. **(A)** Schematic illustration of the in_(C_4_G_2_)75 repeat construct including 6× stop codons, 450bp of human intronic sequences at the N-terminus and 3× protein tags at the C-terminus of the repeats to monitor the DPR proteins in each frame. **(B)** Representative images of antisense RNA foci in HEK293T cells and in primary cortical neurons expressing in_(C_4_G_2_)75 repeats detected by RNA FISH. Red, foci; blue, DAPI; magenta, MAP2. **(C)** Kaplan-Meier curves showing increased risk of cell death in in_(C_4_G_2_)75 expressing primary cortical neurons compared with 2 repeats. Statistical analyses were performed using Mantel-Cox test. **(D-E)** Immunoblotting analysis of phosphorylated PKR (p-PKR) and total PKR in HEK293T cells expressing in_(C_4_G_2_)75 or 2 repeats. p-PKR levels were detected using anti-p-PKR (T446) and normalized to total PKR. GAPDH was used as a loading control. Error bars represent S.D. (n=3 independent experiments). Statistical analyses were performed using student’s t-test. **(F-G)** Immunoblotting analysis of Phosphorylated eIF2α (p-eIF2α) and total eIF2α in HEK293T cells expressing in_(C_4_G_2_)75 or 2 repeats. p-eIF2α levels were detected using anti-phosphor eIF2α (Ser51) and normalized to total eIF2α. GAPDH was used as a loading control. Error bars represent S.D. (n=3 independent experiments). Statistical analyses were performed using student’s t-test. (**H)** Immunoblotting analysis of p-PKR and p-eIF2α in HEK293T cells expressing in_(C_4_G_2_)75, with or without co-expression of wild type PKR, or treatment of a PKR inhibitor, C16. Error bars represent S.D. (n=2 independent experiments). Statistical analyses were performed using one-way ANOVA with Tukey’s post hoc test.

To determine if C9orf72 antisense C_4_G_2_ expanded repeats can cause neuronal toxicity, we co-transfected in_(C_4_G_2_)75 or control 2 repeats together with mApple in mouse primary cortical neurons at 4 days in vitro (DIV4) and used automated longitudinal microscopy to track the survival of hundreds of neurons as indicated by the mApple fluorescence over days. Neurons expressing in_(C_4_G_2_)75 die much faster than those expressing control 2 repeats, suggesting that C9rof72 antisense C_4_G_2_ expanded repeats are neurotoxic (**Fig. 1C**).

### C9orf72 antisense C_4_G_2_ expanded repeats activate PKR/eIF2α-dependent integrated stress response

We next investigated the molecular mechanism underlying toxicity caused by C9orf72 antisense C_4_G_2_ expanded repeats. More than 50 neurological diseases are genetically associated with microsatellite repeat expansions. Repeat expanded RNAs, including CAG, CUG, CCUG, CAGG, and G_4_C_2_, have been shown to activate the double-stranded RNA-dependent protein kinase (PKR) [61, 62]. We hypothesized that C9orf72 antisense C_4_G_2_ expanded RNAs can also activate PKR. HEK293T cells expressing in_(C_4_G_2_)75 show a significant increase in the level of phosphorylated PKR compared to cells expressing 2 repeats, while the total level of PKR remains unchanged (**Fig. 1D-E)**. PKR is one of four kinases that are activated during the integrated stress response (ISR), an evolutionarily conserved stress signaling pathway that adjusts cellular biosynthetic capacity according to need. The four ISR kinases, including PKR, PKR-like ER kinase (PERK), heme-regulated eIF2α kinase (HRI) and general control non-derepressible 2 (GCN2), respond to distinct environmental and physiological stresses by phosphorylating the eukaryotic translation initiation factor eIF2α to cause a temporary shutdown of global protein synthesis and upregulation of specific stress-responsive genes [63]. Accompanying PKR activation, in_(C_4_G_2_)75 significantly increases the phosphorylation of eIF2α without affecting its total level (**Fig. 1F-G**). in_(C_4_G_2_)75 activates eIF2α mainly by the phosphorylation of PKR as other IRS kinases such as PERK phosphorylation are not altered (**Fig. S2A**). Consistent with this, overexpressing wild type (WT) PKR further increases the phosphorylation of eIF2α induced by in_(C_4_G_2_)75, whereas treatment with a specific PKR inhibitor C16 reduces the phosphorylation of both PKR and eIF2α to a level comparable to that of cells expressing 2 repeats **(Fig 1H-I)**. We further expressed in_(C_4_G_2_)75 in a neuronal cell line SH-SY5Y that is commonly used to study neurodegeneration and observed similar activation of PKR and eIF2α by the antisense expanded repeats (**Fig. S2B-C**).

To determine whether the activation of PKR/eIF2α leads to a global mRNA translation inhibition, we employed a puromycin-based, nonradioactive method to monitor protein synthesis [64]. Puromycin is a structure analog of aminoacyl-tRNA that incorporates into nascent polypeptide chains and prevents elongation. The amount of incorporated puromycin detected by antibodies reflects global translation efficacy. HEK293T cells expressing in_(C_4_G_2_)75 show a significantly reduced amount of incorporated puromycin compared to those expressing 2 repeats (**Figs. 2A and S2D**). Similarly, neurons expressing 75 antisense repeats, as identified by GP DPR accumulation, have robust global translation inhibition (**Fig. 2B**). In response to stress-induced translation inhibition, we also observed abundant accumulation of stress granules. Approximately 32% of cells expressing in_(C_4_G_2_)75, identified by the presence of antisense RNA foci, show G3BP1-positive stress granules, and ∼55% of foci-positive cells stain for FMRP, another commonly used marker for stress granules **(Fig. 2C-D)**. These results support that C9orf72 antisense C_4_G_2_ expanded repeats activate PKR/eIF2α-dependent integrated stress response, leading to global translation inhibition and stress granule formation.

**Figure 2.**
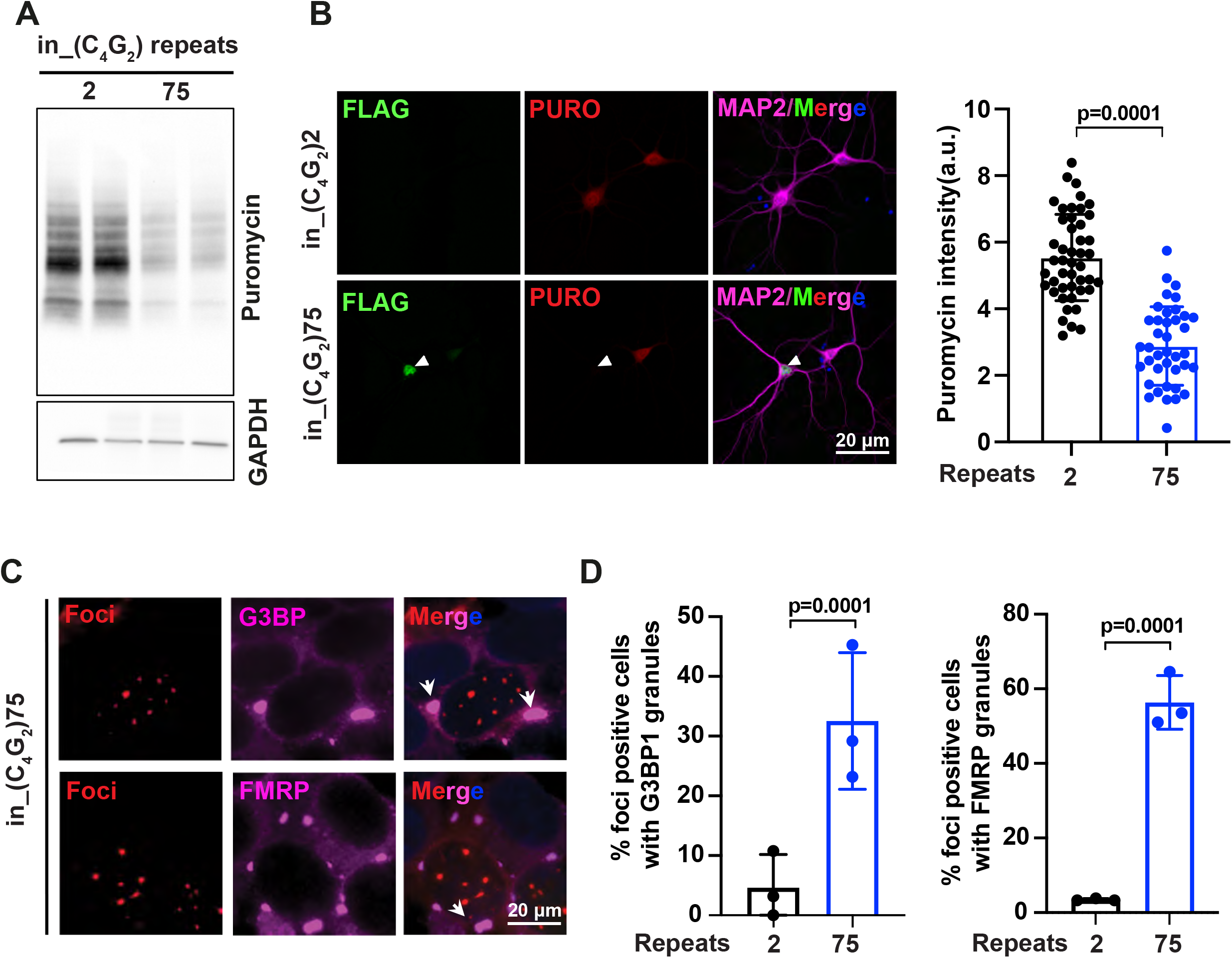
C9orf72 C_4_G_2_expanded repeats inhibit global protein synthesis and induce stress granule assembly. **(A)** Immunoblotting of puromycin in HEK293T cells expressing in_(C_4_G_2_)75 or 2 repeats. Cells were incubated with puromycin for 30 minutes before harvesting. **(B)** Representative images of primary neurons expressing either (C_4_G_2_)75 or 2 repeats stained with anti-puromycin (red), anti-FLAG (green), DAPI (blue) and MAP2 (magenta). Quantification of puromycin intensity in neurons expressing in_(C_4_G_2_)75 or 2 repeats. Error bars represent S.D. (n=40-50 neurons/group. Similar results were obtained from two independent experiments). Statistical analyses were performed using student’s t-test. **(C)** Representative images of G3BP1 and FMRP staining in HEK293T cells expressing in_(C_4_G_2_)75 repeats identified by the presence of RNA foci using FISH. **(D)** Quantification of antisense foci positive cells with G3BP1 and FMRP granules. Error bars represent S.D. (n=150 cells/condition and three independent experiments). Statistical analyses were performed using student’s t-test.

### Antisense C9orf72 repeat expanded RNAs activate the PKR/eIF2α pathway independent of DPR proteins

We next determined whether the activation of PKR/eIF2α-dependent integrated stress response is driven by repeat RNA themselves and/or by dipeptide repeat proteins. We first expressed individual codon-optimized, ATG-driven DPR proteins. Neither PR50, PA50 or GP80 activate the phosphorylation of eIF2α, suggesting that the activation of the PKR/eIF2α pathway by C9orf72 antisense C_4_G_2_ expanded repeats is unlikely due to the DPR proteins produced by RAN translation (**Fig. S3A-B**). To obtain direct evidence that C9orf72 antisense repeat expanded RNAs activate PKR/eIF2α themselves, we used two strategies to reduce/inhibit DPR proteins without affecting the RNA. First, recent studies have shown that C9orf72 G_4_C_2_ sense repeat expanded RNAs initiate RAN translation at a near-cognate CUG codon in the intronic region 24 nucleotides upstream of the repeat sequence [9, 65]. We thus hypothesized that RNA translation from C_4_G_2_ antisense repeat expanded RNAs might similarly depend on the intronic sequence at the 5’ region. We generated a new construct (C_4_G_2_)75 without including the 450bp human intronic sequence **(Fig. 3A)**. Supporting our hypothesis, cells expressing (C_4_G_2_)75 do not accumulate any detectable GP, PA, or PR DPR proteins, which is strikingly different compared to those expressing in_(C_4_G_2_)75 with the 450bp human intronic sequence **(Fig. 3B)**. The reduced/abolished DPR proteins by (C_4_G_2_)75 are not due to altered RNA expressions since levels of RNA transcripts and antisense foci are comparable to those of in_(C_4_G_2_)75 **(Figs. 3C and S3C)**. Second, we obtained a previously reported stop codon-interrupted 108 antisense repeat construct, designated as RNA only (RO) [(C_4_G_2_)108RO]. It has been shown that this construct is not RAN translated to produce DPR proteins, while still adopts similar stable conformations as the uninterrupted repeat RNAs [51, 52]. As expected, no detectable antisense DPR proteins are observed in cells expressing (C_4_G_2_)108RO, despite abundant accumulation of antisense foci (**Fig. S3C-D**). Interestingly, expression of either (C_4_G_2_)75 or (C_4_G_2_)108RO leads to the robust activation of PKR and eIF2α at a comparable level as seen for in_(C_4_G_2_)75 (**Fig. 3D-E)**. Thus, C9orf72 antisense repeat expanded RNAs activate the PKR/eIF2α pathway independent of DPR proteins.

**Figure 3.**
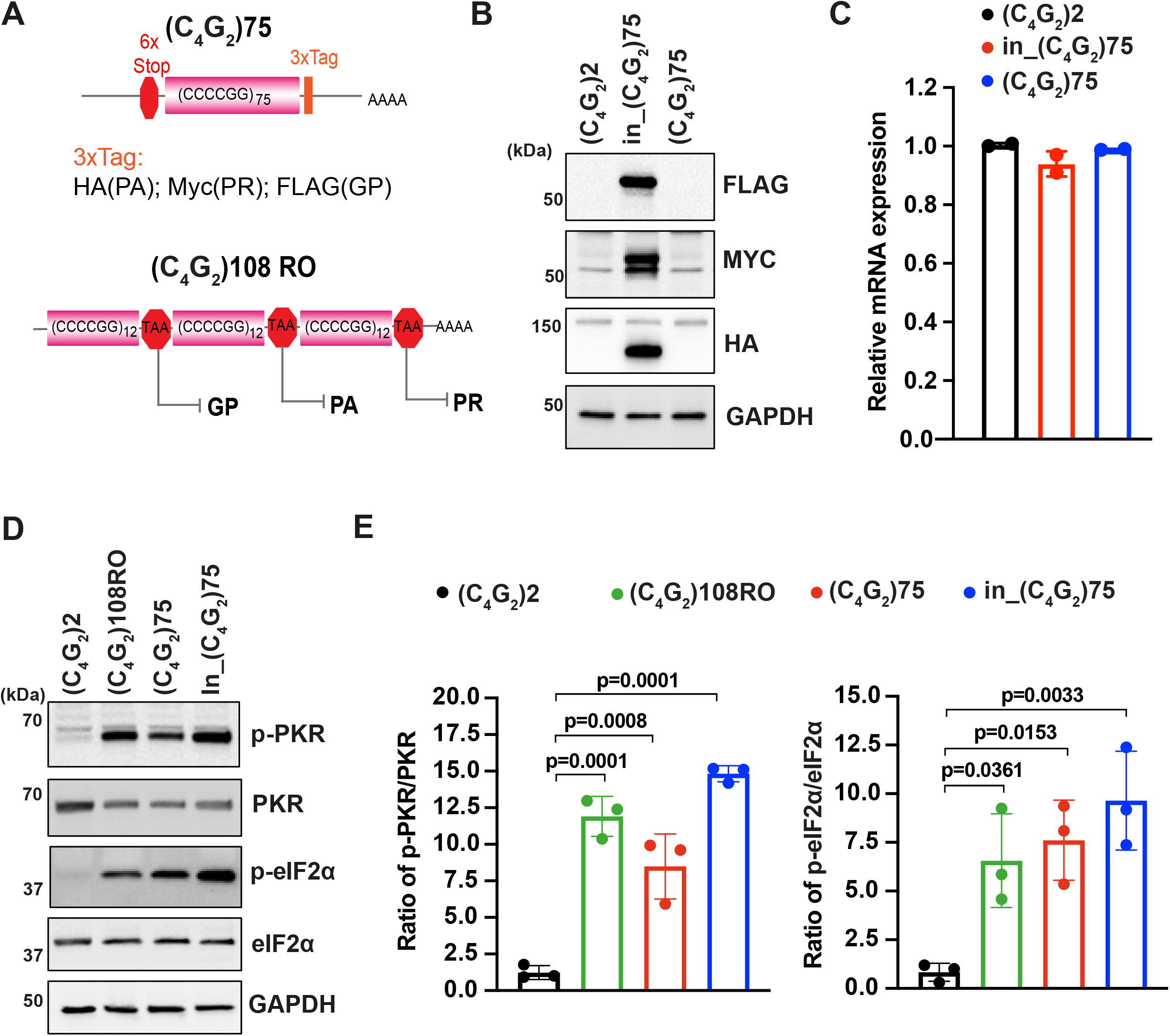
Antisense C9orf72 repeat expanded RNAs activate the PKR/eIF2α pathway independent of DPR proteins. **(A)** (Top) Schematic illustration of (C_4_G_2_)75 repeats without the human intronic sequences. 3× protein tags were included at the C-terminus of the repeats to monitor the DPR proteins in each frame. (Bottom) Schematic illustration of antisense (C_4_G_2_)108RO repeats with stop codons inserted in every 12 repeats to prevent the translation of DPR proteins from all reading frames. **(B)** Immunoblotting of DPR proteins in HEK293T cells expressing in_(C_4_G_2_)75, (C_4_G_2_)75, or 2 repeats. DPR protein levels were detected using anti-FLAG (frame with GP), anti-MYC (frame with PR), and anti-HA (frame with PA). GAPDH was used as a loading control. **(C)** mRNA levels were measured by quantitative qPCR in cell expressing in_(C_4_G_2_)75, (C_4_G_2_)75, or 2 repeats. Error bars represent S.D. (n=2). **(D-E)** Immunoblotting of p-PKR and p-eIF2α in HEK293T cells expressing in_(C_4_G_2_)75, (C_4_G_2_)75, or 2 repeats. p-PKR (T446) and p-eIF2α (Ser51) were normalized to total PKR and eIF2α respectively. GAPDH was used as a loading control. Error bars represent S.D. (n=3 independent experiments). Statistical analyses were performed using one-way ANOVA with Tukey’s post hoc test.

### Antisense C9orf72 repeat expanded RNAs themselves induce stress granules and lead to neuronal toxicity

Given the conflicting reports of whether C9orf72 antisense RNAs themselves are toxic independent of DPR proteins [11, 51, 52], we next focused on (C_4_G_2_)108RO, which is capable of activating PKR/eIF2α. We first determined whether this is sufficient to induce stress granules. FMRP is diffused in the cytoplasm of cells expressing 2 C_4_G_2_ repeats as seen before, but it rapidly assembles into stress granules in cells expressing (C_4_G_2_)108RO. This suggests that antisense C_4_G_2_ repeat expanded RNAs themselves can trigger stress granule formation in the absence of DPR proteins. (**Fig. 4A-B**). To determine the role of PKR activation in stress granule formation by (C_4_G_2_)108RO, we knocked down PKR using siRNAs. siRNAs targeting PKR reduce its protein level by 80% compared to control siRNAs (**Fig. 4C and S3E**). Consequently, the phosphorylation of eIF2α by (C_4_G_2_)108RO is almost inhibited (**Fig. 4C and S3E)** and the percentage of foci-positive cells with stress granules is significantly reduced (**Fig. 4D-E**). This data suggests that C9orf72 antisense repeat expanded RNAs themselves induce stress granules by activating PKR/eIF2α.

**Figure 4.**
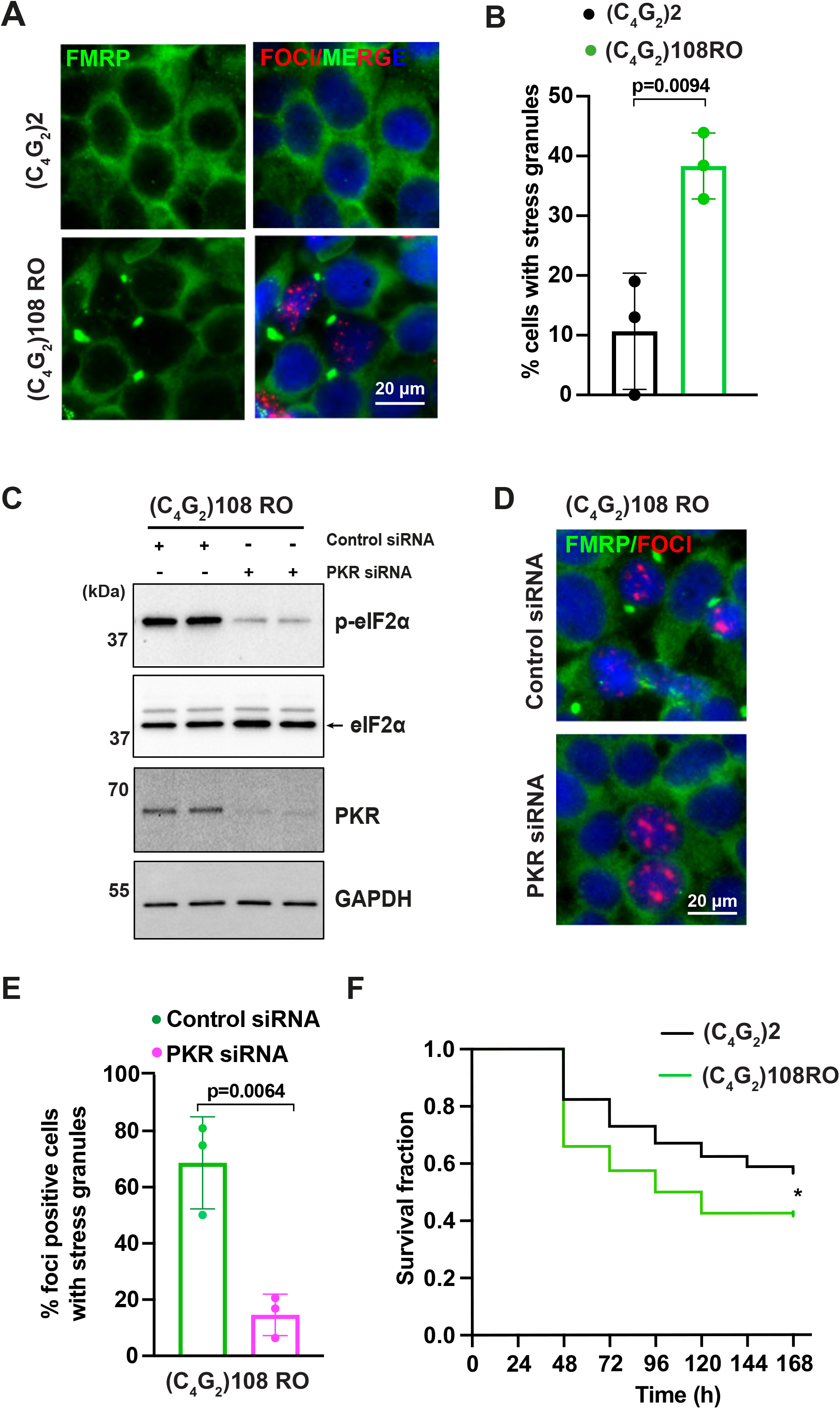
Antisense C9orf72 repeat expanded RNAs themselves induce stress granules and lead to neuronal toxicity. **(A)** Representative images of FMRP staining in HEK293T cells expressing (C_4_G_2_)108RO or 2 repeats. **(B)** Quantification of antisense foci positive cells with FMRP granules. Error bars represent S.D. (n=180 cells/condition and three independent experiments). Statistical analyses were performed using student’s t-test. **(C)** Immunoblotting of PKR and p-eIF2α (Ser51) in HEK293T cells expressing (C_4_G_2_)108RO together with control or PKR siRNA. GAPDH was used as a loading control. **(D-E)** Representative images of FMRP staining in HEK293T cells expressing (C_4_G_2_)108 repeats together with either control or PKR siRNA **(D)**. Quantification of antisense foci positive cells with FMRP granules. Error bars represent S.D. (n=150 cells/condition and three independent experiments) **(E). (F)** Kaplan-Meier curves showing increased risk of cell death in (C_4_G_2_)108RO expressing neurons compared with 2 repeats. Statistical analyses were performed using Mantel-Cox test (* p<0.05).

We further utilized the unbiased longitudinal microscopy assay to determine the risk of death in neurons expressing (C_4_G_2_)108RO. Rodent primary cortical neurons were transfected with mApple and (C_4_G_2_)108RO or 2 repeats and imaged at 24-hour intervals for 10 days. Neurons expressing (C_4_G_2_)108RO show a significant decrease in survival compared to control neurons expressing 2 repeats, suggesting that C9orf72 antisense repeat expanded RNAs themselves are neurotoxic (**Fig. 4F**).

### Increased levels of phosphorylated PKR and eIF2α in C9FTD/ALS patients

To study disease relevance, we determined the levels of phosphorylated PKR and eIF2α in C9FTD/ALS patient postmortem tissues. Immunohistochemistry staining showed that the level of phosphorylated PKR is increased in the frontal cortex, especially in the large pyramidal neurons, of patients carrying C9orf72 repeat expansions compared to age-matched non-disease controls (**Fig. 5A**). In addition, the level of phosphorylated eIF2α is also significantly increased after normalizing to the total eIF2α level, despite the heterogeneity of eIF2α protein levels in patients (**Fig. 5B**). These results suggest that the PKR/eIF2α pathway is activated in C9FTD/ALS patients.

**Figure 5.**
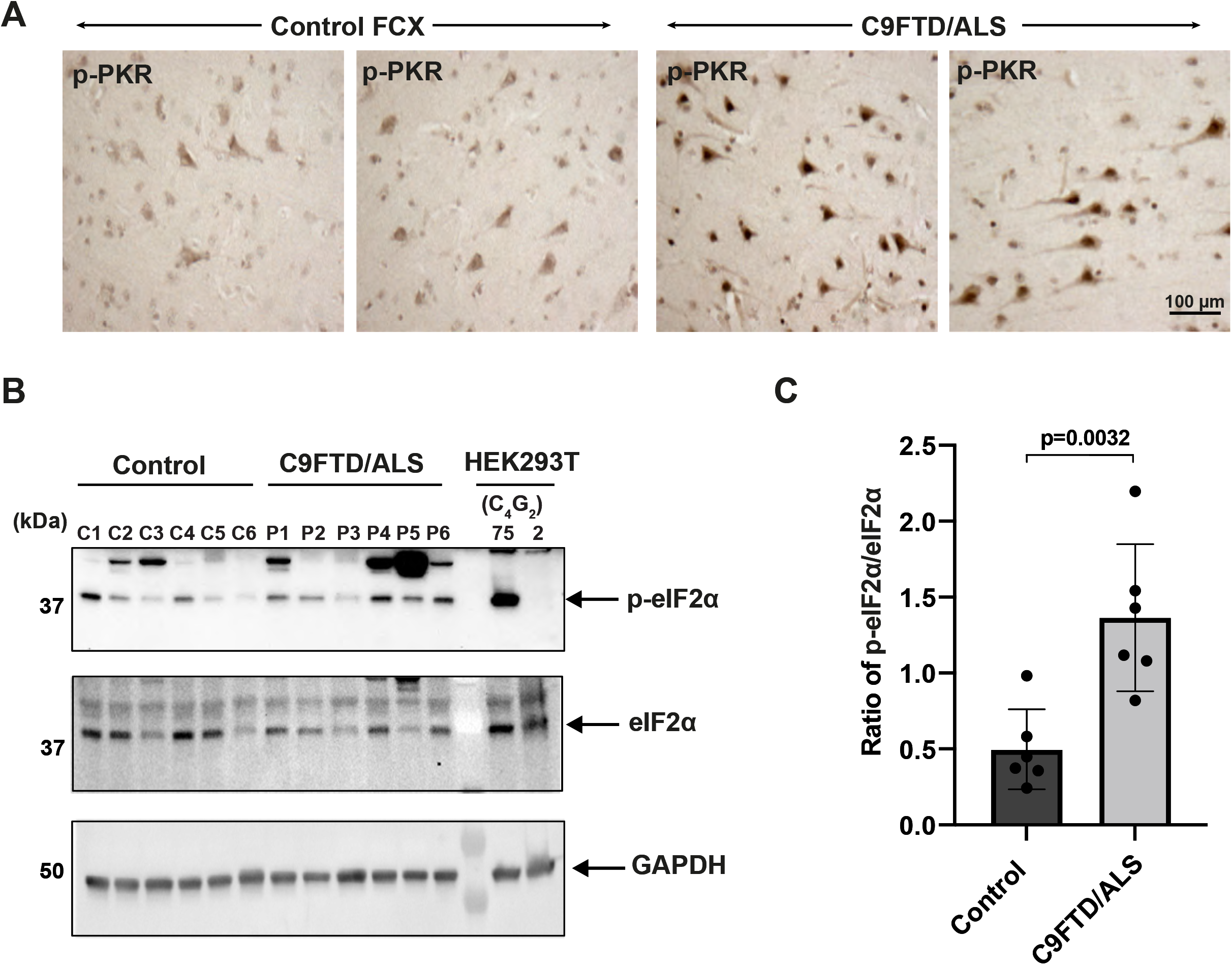
Increased levels of phosphorylated PKR and eIF2α in C9FTD/ALS patients. **(A)** Representative immunohistochemistry images of phosphorylated PKR staining in control and C9FTD/ALS patient’s frontal cortex (FCX) using anti-p-PKR (T446) (n=4 per genotype). (**B-C)** Immunoblotting of p-eIF2α in proteins extracted from control (C1-C6) and C9FTD/ALS patient’s frontal cortex (P1-P6). p-eIF2α (Ser51) was normalized to total eIF2α. GAPDH was used as a loading control. Error bars represent S.D. (control n=6 and C9FTD/ALS n=6). Statistical analyses were performed using unpaired student’s t-test.

### Sense C9orf72 repeat expanded RNAs cannot activate the PKR/eIF2α pathway

By expressing a construct containing (G_4_C_2_)120, Zu *et al*. showed that this repeat expansion construct activates PKR and increases DPR protein translation in HEK293T cells [66]. However, it is unknown whether this construct produces antisense (C_4_G_2_) transcripts that are responsible for the PKR activation. Therefore, we generated a construct with similar repeat length, (G_4_C_2_)75 (**Fig. S4A**). Consistent with the earlier findings by Zu *et al*., expression of (G_4_C_2_)75 in HEK293T cells significantly increases phosphorylation of both PKR and eIF2α (**Fig. S4B-C**). Interestingly, we detected abundant accumulation of both sense and antisense RNA foci in cells expressing (G_4_C_2_)75 but not in those expressing 2 repeats (**Fig. S4D**). To determine the relative contribution of sense (G_4_C_2_) and antisense (C_4_G_2_) repeat expanded RNAs, we first used previously published antisense oligonucleotides (ASOs) that specifically degrade sense (G_4_C_2_) RNAs [22]. As expected, ASOs targeting sense RNA repeats significantly reduce the accumulation of sense RNA foci but have little effect on antisense RNA foci (**Figs. 6A-B and S4E**). However, reducing sense RNA transcripts/foci does not alter the activation of PKR/eIF2α by (G_4_C_2_)75 (**Figs. 6C-D and S4F-G**). We next designed two ASOs specifically targeting antisense repeat RNAs. Both ASO1 and ASO2 targeting C9orf72 antisense RNAs significantly reduce the abundance of antisense RNA foci without affecting sense RNA foci (**Fig. 6E-F**). Consequently, both ASOs significantly inhibit the activation of PKR and eIF2α by (G_4_C_2_)75 (**Fig. 6G-H**). Thus, antisense (C_4_G_2_), but not sense (G_4_C_2_), C9orf72 repeat expanded RNAs activate the PKR/eIF2α pathway.

**Figure 6.**
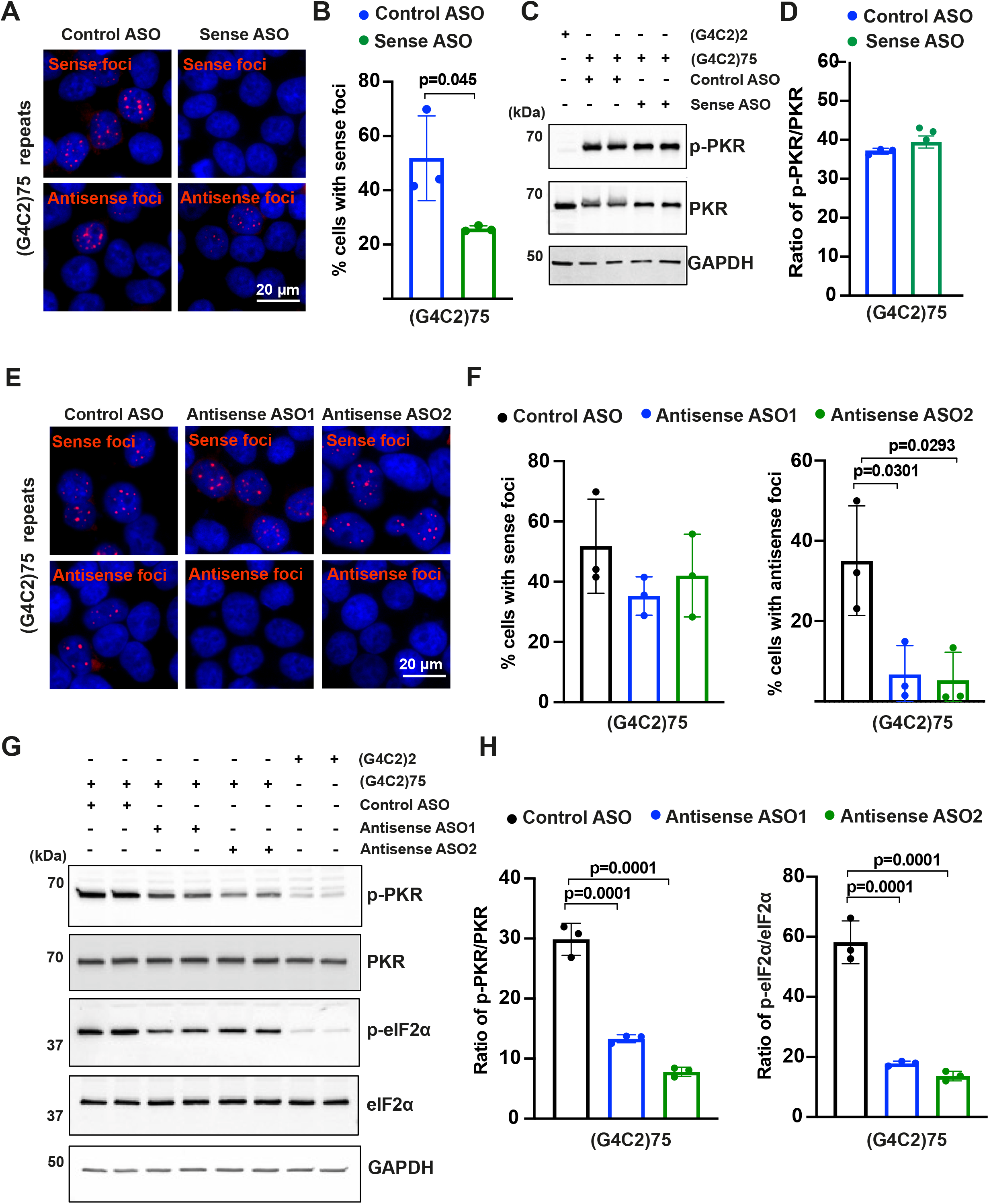
Sense C9orf72 repeat expanded RNAs cannot activate the PKR/eIF2α pathway. **(A-B)** Representative images **(A)** and Quantification of sense and antisense RNA foci in HEK293T expressing (G_4_C_2_)75 repeats together with either control ASOs or ASOs targeting sense (G_4_C_2_) repeat expanded RNAs (**B)**. Foci were detected by RNA FISH. Red, foci; blue, DAPI. Error bars represent S.D. (n=90 cells/condition from three independent experiments). Statistical analyses were performed using student’s t-test. **(C-D)** Immunoblotting of p-PKR in HEK293T cells expressing (G_4_C_2_)75 or (G_4_C_2_)2 repeats together with either control ASO or ASOs targeting sense (G_4_C_2_) repeat expanded RNAs. Phosphorylated PKR levels were detected using anti-p-PKR (phosphor T446) and normalized to total PKR. GAPDH was used as a loading control. Error bars represent S.D. (n=3 independent experiments). Statistical analyses were performed using unpaired student’s t-test. **(E-F)** Representative images **(E)** and Quantification **(F)** of sense and antisense RNA foci in HEK293T expressing (G_4_C_2_)75 repeats together with either control ASOs or ASOs targeting sense repeat expanded RNAs. Foci were detected by RNA FISH. Red, foci; blue, DAPI. Error bars represent S.D. (n=90 cells/condition and 3 independent experiments). Statistical analyses were performed using student’s t-test. (**G-H)** Immunoblotting of p-PKR and p-eIF2α in HEK293T cells expressing (G_4_C_2_)75 or (G_4_C_2_)2 repeats together with either control ASO or ASOs targeting antisense (G_4_C_2_) repeat expanded RNAs **(G)**. P-PKR (T446) and p-eIF2α (Ser51) were normalized to total PKR and eIF2α respectively **(H)**. GAPDH was used as a loading control. Error bars represent S.D. (n=3 independent experiments). Statistical analyses were performed using one-way ANOVA with Tukey’s post hoc test.

## Discussion

It is generally accepted in the field that the bidirectionally transcribed repeat expanded RNAs play important roles in FTD/ALS caused by C9orf72 repeat expansions [4]. However, the relative contributions of potential toxic species, including sense and antisense RNAs themselves, RNA foci, and DPR proteins, are largely debated. Our study shows for the first time that C9orf72 antisense (C_4_G_2_), but not sense (G_4_C_2_), repeat expanded RNAs activate PKR/eIF2α-dependent integrated stress response and lead to neurotoxicity independent of DPR proteins in model systems. We also detected increased activation of PKR/eIF2α in the frontal cortex of C9FTD/ALS patients. Consistent with our observations, increased phosphorylation of PKR has also been reported in BAC transgenic mice expressing 500 G_4_C_2_ repeats and in C9orf72 patients by two other studies [67, 68].

Several studies argue against the toxicity from sense (G_4_C_2_) repeat expanded RNAs and associated RNA foci. In one study, *Drosophila* expressing intronic (G_4_C_2_)160 repeats show abundant sense RNA foci in the nucleus but have little DPR proteins and no neurodegeneration, suggesting that sense RNA foci is insufficient to cause toxicity in this model [52]. Two other elegant studies generated *Drosophila* expressing interrupted (G_4_C_2_) repeats by inserting stop codons every 12 repeats in all reading frames to prevent RAN translation. These *Drosophila* do not show any toxicity whereas those expressing pure (G_4_C_2_) repeats of similar sizes do, despite comparable sense RNA foci accumulation. Similarly, *Drosophila* expressing interrupted antisense repeat expanded RNAs do not show any deficits, suggesting that antisense (C_4_G_2_) RNAs are not toxic in this model system. However, expressing the same antisense interrupted repeat construct (C_4_G_2_)108RO causes motor axonopathy in zebrafish [11]. Consistent with this, we show that (C_4_G_2_)108RO is also toxic to primary cortical neurons. It is interesting to note that PKR, which is constitutively and ubiquitously expressed in vertebrate cells including zebrafish, is not found in plants, fungi, protists, or invertebrates such as *Drosophila* [63].

The contribution of antisense repeat expanded RNAs to C9FTD/ALS pathogenesis is understudied, although PR DPR proteins RAN translated from the antisense RNAs have been shown to be toxic in various model systems [25, 36, 69]. However, aggregates of PR DPR proteins are rare in C9FTD/ALS postmortem tissues, whereas antisense RNA foci are as abundant as sense RNA foci in multiple CNS regions despite scarcity of antisense RNA transcripts. One possibility of such discrepancy between RNA transcript and foci levels is that antisense RNA foci are extraordinarily stable and rarely turn over once formed along the lifespan of patients. Several neuropathological studies have attempted to correlate the abundance and distribution of antisense RNA foci with C9FTD/ALS clinical features. Mizielinska *et al*. showed that patients with more antisense RNA foci tend to have an earlier age of symptom onset [54] and more intriguingly, antisense RNA foci are shown to be associated with nucleoli and mislocalization of TDP-43 in two different studies [70, 71]. These results highlight the disease relevance of antisense repeat expanded RNAs in C9FTD/ALS and the significance of our work. Our study, however, does not differentiate antisense RNA foci from RNAs themselves since it is technically challenging given that RNA foci inevitably form with the expression of repeat expanded RNAs. The proposed mechanisms of RNA foci-mediated toxicity are via sequestration of critical RBPs. Several RBPs have been proposed to interact with and/or are sequestered into sense RNA foci, yet those interacting with antisense RNA foci have not been well characterized and are worth exploring especially in correlation with PKR/eIF2α activation.

How is PKR specifically activated by C9orf72 antisense, but not sense repeat expanded RNAs? PKR is a stress sensor first identified as a kinase responding to viral infections by directly binding to viral double stranded RNAs (dsRNAs) [63]. Several disease relevant repeats expanded RNAs, such as CUG and CGG, have been shown to form stable hairpins and directly bind to PKR, leading to its activation [61, 62]. It is possible that antisense (C_4_G_2_) RNAs form similar hairpin structures, which has not been well studied. Supporting this, antisense (C_4_G_2_) DNAs form i-motifs consisting of two parallel duplexes in a head to tail orientation as well as protonated hairpins under near-physiological conditions [72]. In contrast, sense (G_4_C_2_) RNAs tend to form stable unimolecular and multimolecular G-quadruplexes [73, 74]. In addition to C9orf72 antisense repeat expanded RNAs, short and long interspersed retrotransposable elements (SINEs and LINEs) and endogenous retroviruses (ERVs) represent other main sources of endogenous dsRNAs [75]. In C9orf72 patients, transcripts from multiple classes of repetitive elements are significantly elevated [76]. Other PKR activators include cellular stresses such as oxidative stress, intracellular calcium increase or ER stress, as well as interferon-gamma (IFNγ), tumor necrosis factor α (TNFα), heparin, and platelet-derived growth factor [63]. Whether C9orf72 antisense (C_4_G_2_) expanded RNAs specifically increase transcription of RNAs with repetitive elements or other PKR activators warrants additional studies.

Our study shows that C9orf72 antisense (C_4_G_2_) expanded repeats promotes robust global translation inhibition and stress granule formation independent of DPR proteins via the activation of the PKR/eIF2α pathway. Stress granules are dynamic structures that form and disperse rapidly with acute stress. However, chronic stress during aging or under pathological conditions leads to altered stress granule dynamics and persistent stress granules, which have been implicated to the aggregation of RBPs such as TDP-43 and contribute to the pathogenesis of FTD and ALS. For C9orf72 FTD/ALS, several studies show that DPR proteins such as GR, PR and GA are toxic [23-25, 36-50]. It has been shown that GR and PR can also inhibit global protein translation via direct binding to mRNAs to block access to the translational machinery [40]. Interestingly, the RAN translation of DPR proteins is specifically increased by the integrated stress response via eIF2α phosphorylation [65, 77]. Thus, activation of PKR/eIF2α by C9orf72 antisense repeat expanded RNAs will lead to additional accumulation of DPR proteins and toxicity.

From the therapeutic development point of view, several approaches have been explored to mitigate gain of toxicity from C9orf72 repeat expanded RNAs [4]. Our previous work with ASOs targeting C9orf72 sense (G_4_C_2_) repeat expanded RNAs showed great promise in a preclinical mouse model expressing 450 (G_4_C_2_) repeats [22]. Unfortunately, there was a recent setback in the clinical trial using these ASOs to treat C9ALS patients. Although many different confounding reasons may cause drug failure, further understanding of disease mechanisms is required to develop successful therapies for C9FTD/ALS. On this note, Zu *et al*. recently showed that metformin, an FDA approved drug widely used for treating type 2 diabetes, inhibits PKR activation, reduces DPR proteins RAN translated from the sense strand, and improves behavioral and pathological deficits in BAC transgenic mice expressing G_4_C_2_ repeats [68]. Our study highly suggests that the activation of PKR in these BAC transgenic mice and in C9orf72 patients might result from the antisense repeat expanded RNAs. Future therapies targeting C9orf72 antisense RNAs and/or altered downstream molecular pathways hold great promise for these devastating neurodegenerative diseases.

## Materials and Methods

### Plasmids and siRNAs

A construct containing 10 GGGGCC repeats, flanked 5’ by BbsI and 3’ by BsmBI recognition sites, was synthesized by GENEWIZ and used to generate antisense (C_4_G_2_) repeats using recursive directional ligation as previously described [24]. The repeat-containing plasmids were amplified using recombination deficient Stbl3 *E. coli* (Life Technologies) at 32°C to minimize retraction of repeats. Human PKR cDNA was a gift from Dr. Thomas Dever (NIH, USA) and (C_4_G_2_)108RO was gifted by Dr. Adrian Isaac [51]. For longitudinal fluorescence microscopy pGW1-mApple was used. All plasmids were verified by Sanger sequencing (Genewiz, USA). All ASOs were synthesized by Integrated DNA Technologies, USA. siRNAs against PKR and control siRNAs were purchased from Horizon Discovery, USA.

### Human tissues

Post-mortem brain tissues from C9FTD/ALS patients (n=6) and controls (n=6) were obtained from the Emory Neuropathology Core. Patient information is provided in Table S1.

### Cell culture and transfection

Human embryonic kidney (HEK293T) and human neuroblastoma (SH-SY5Y) cells from ATCC were cultured in high glucose DMEM (Invitrogen) and DMEM-F12 (Invitrogen), respectively (supplemented with 10% fetal bovine serum (Corning), 4 mM Glutamax (Invitrogen), penicillin (100 U/mL), streptomycin (100 μg/mL) and non-essential amino acids (1%). Cells were grown at 37°C in a humidified atmosphere with 5% CO_2_. Cells were transiently transfected using polyethyleneimine or lipofectamine. Experiments were performed 48 hours after transfection.

### Primary cortical neuronal culture and transfection

Primary cortical neurons were prepared from C57BL/6J mouse embryos (Charles River) of either sex on embryonic day 17. Cerebral cortices were dissected and enzymatically dissociated using trypsin with EDTA (Thermo Fisher Scientific; 10 minutes), mechanically dissociated in Minimum Essential Media (MEM; Fisher) supplemented with 0.6% glucose (Sigma) and 10% Fetal Bovine Serum (FBS; Hyclone) and stained to assess viability using Trypan Blue (Sigma). Neurons were plated on coverslips (Matsunami Inc., 22 mm) or MatTek dishes coated with poly-l-lysine (Sigma). A total of 50,000 neurons were plated as a ‘spot’ on the center of the coverslip to create a small, high-density network. Neurons were cultured in standard growth medium [glial conditioned neurobasal plus medium (Fisher) supplemented with Glutamax (GIBCO) and B27 plus (Invitrogen)], and half of the media was exchanged 2-3 times a week until the experiment endpoints. No antibiotics or antimycotics were used. Cultures were maintained in an incubator regulated at 37 °C, 5% CO_2_ and 95% relative humidity as described [78]. Cells were transiently transfected using Lipofectamine 2000 (Invitrogen) according to the manufacturer’s instructions.

### Longitudinal fluorescence microscopy

Mouse primary cortical neurons were transfected with mApple and repeat expanded constructs and imaged by fluorescence microscopy at 24-hour intervals for 7-10 days as described [79]. Time of death was determined based on rounding of the soma, retraction of neurites, or loss of fluorescence. The time of death for individual neurons was used to calculate the risk of death in each population relative to a reference group. Images were acquired using Keyence BZ-X810 microscope with a 10× objective and analyzed by Image J. The images were stitched and stacked, and cell death was scored using the criteria mentioned above.

### RNA fluorescence in situ hybridization

LNA DNA probes were used against the sense and antisense hexanucleotide repeat expanded RNAs (Exiqon, Inc.). The probe sequence for detecting sense RNA foci: TYE563-CCCCGGCCCCGGCCCC; and that for antisense RNA foci is: TYE563-GGGGCCGGGGCCGGGG. All hybridization steps were performed under RNase-free conditions. Cells were fixed in 4% paraformaldehyde (Electron Microscopy Sciences) for 20 minutes, washed three times for 5 minutes with phosphate buffer saline (DEPC 1× PBS, Corning) followed by permeabilization with 0.2% Triton-X 100 (Sigma) for 10 minutes and then incubated with 2× SSC buffer for 10 minutes. Cells were hybridized (50% formamide, 2× SCC, 50 mM sodium phosphate (pH 7), 10% dextran sulfate, and 2 mM vanadyl sulfate ribonucleosides) with denatured probes (final concentration of 40 nM) at 66°C for 2 hours. After hybridization, slides were washed at room temperature in 0.1% Tween-20/2×SCC for 10 minutes twice and in stringency washes in 0.1× SCC at 65°C for 10 minutes. Cell nuclei were stained with DAPI. Three to six random pictures were taken by Keyence BZ-X810 microscope with a 60× oil objective and analyzed by Image J.

### Immunofluorescence

Cells were fixed in 4% paraformaldehyde (Electron Microscopy Sciences) for 20 minutes, washed three times for 5 minutes with phosphate buffer saline (1× PBS, Corning) and treated with 0.2% Triton-X 100 (Sigma) in PBS for 10 minutes. Cells were blocked for 30 minutes in a blocking solution consisting of 4% bovine serum albumin (Sigma) in PBS. Cells were incubated overnight in primary antibodies diluted in blocking solution. The next day, cells were washed 3 times for 5 minutes in PBS and incubated in secondary antibodies in blocking solution for one hour at room temperature (dark). After washing 3 times for 5 minutes, coverslips with the cells were mounted using Prolong Gold Antifade mounting media (Invitrogen). Images were acquired with Keyence BZ-X810 microscope with a 60× oil objective and analyzed by Image J.

### Immunohistochemistry

Post-mortem brain tissues were obtained from the brain bank maintained by the Emory Alzheimer Disease Research Center under proper Institutional Review Board protocols. Paraffin-embedded sections from frontal cortex (8 μm thickness) were deparaffinized by incubation at 60°C for 30 minutes and rehydrated by immersion in graded ethanol solutions. Antigen retrieval was done by microwaving in a 10 mM citrate buffer (pH 6.0) for 5 minutes followed by allowing slides to cool to room temperature for 30 minutes. Endogenous peroxidase activity was eliminated by incubating slides with hydrogen peroxide block solution (Fisher) for 10 minutes at room temperature followed by rinsing in phosphate buffered saline. Non-specific binding was reduced by blocking in ultra-Vision Block (Fisher) for 5 minutes at room temperature. Sections were then incubated overnight with primary antibodies diluted in 1% BSA in phosphate buffered saline for 30 minutes at room temperature or incubated without primary antibody as a negative control. Sections were rinsed in phosphate buffered saline and incubated in labeled ultra Vision LP detection system horseradish peroxidase-polymer secondary antibody (Fisher) for 15 minutes at room temperature. Slides were imaged for analysis using an Aperio Digital Pathology Slide Scanner (Leica Biosystems). For IHC, rabbit anti-p-PKR, Millipore 07-532 (1:100 dilution) antibody was used.

### Protein lysate preparation

Whole cell/tissue extracts were lysed using RIPA Lysis Buffer pH 7.4 (Bio-world, USA) supplemented with Halt™ protease and phosphatase inhibitor cocktail (ThermoFisher Scientific). Lysates were sonicated at 25% amplitude for 3 cycles for 15 seconds with 5 second intervals. Supernatant was collected after centrifuging at max speed for 15 minutes at 4°C. The concentration of the isolated proteins was determined using BCA Protein Assay Reagent (Pierce, USA).

### Immunoblotting assay

For western blotting, 20-30 µg of proteins were prepared in 4× laemmli sample buffer and heat-denatured at 95°C for 5 minutes. Samples were resolved on 4–20% gradient gels (Bio-Rad). Proteins were transferred to nitrocellulose membranes (0.2 µm, Bio-Rad). The membrane was blocked in 5 % milk and incubated overnight at 4°C with primary antibodies diluted in blocking buffer. Secondary antibodies HRP-conjugated secondary antibodies (Abclonal) or IRDye secondary antibodies (Li-cor) were diluted in blocking buffer and applied to the membrane for 1 hour at room temperature. Primary antibodies used: mouse anti-FLAG (1:1000; Sigma), rabbit anti-HA (1:1000; CST), mouse anti-MYC (1:1000; Sigma), rabbit anti-PKR (1:1000; abcam), rabbit anti-phospho-PKR (1:1000; abcam), rabbit anti-eIF2α (1:1000; CST), rabbit anti-phospho-eIF2α (1:1000; CST), rabbit anti-PERK (1:1000; CST), rabbit anti-phospho-PERK (1:1000; abcam), rabbit anti-GAPDH (1:5000; CST). Antibodies against PR, GP and PA have been previously reported [22]. Super Signal West Pico (Pierce, USA) was used for detection of peroxidase activity. Molecular masses were determined by comparison to protein standards (Thermo Scientific). The immunoreactive bands were detected by ChemiDoc Image System (Bio-Rad, USA).

### Quantitative real-time PCR

Total RNAs were extracted using a RNeasy kit as instructed by the manufacturer (Qiagen). cDNA was prepared using High-Capacity cDNA Reverse Transcription Kit from applied biosystem. Quantitative RT-PCR reactions were conducted and analyzed on a StepOnePlus Real-Time PCR system (Applied Biosystems). Gene expression levels were measured by SYBR green (Thermo Fisher Scientific) quantitative real-time PCR.

### Statistical analysis

Statistical analyses and graphs were prepared in GraphPad Prism (version 9). Data is expressed as mean ± S.D. as shown in figure legends. Student t-test or one-way ANOVA was used for statistical analysis unless specified in figure legends.

**Table 1.** List of controls (C1-C6) and C9FTD/ALS patients (P1-P6) post-mortem tissues used in this study.

| ID | Primary Neuropathologic Diagnosis | Age at Onset | Age at Death | Sex |
| --- | --- | --- | --- | --- |
| C1 | Control | - | 53 | F |
| C2 | Control | - | 77 | NA |
| C3 | Control | - | 43 | F |
| C4 | Control | - | 57 | M |
| C5 | Control | - | 72 | F |
| C6 | Control | - | 57 | F |
| P1 | FTLD (C9 expansion) | 58 | 71 | M |
| P2 | ALS (C9 expansion) | 54 | 57 | F |
| P3 | FTLD (C9 expansion) | 57 | 66 | M |
| P4 | FTLD (C9 expansion) | 58 | 67 | M |
| P5 | FTLD (C9 expansion) | 62 | 66 | M |
| P6 | ALS (C9 expansion) | - | 69 | NA |

## Acknowledgements

We thank current and past members of Jiang, Bassell labs and Dr. Homa Ghalei for many helpful discussions. We would like to thank Dr. Jonathan Glass for providing access to the patient samples (Emory cohort). We would like to thank Dr. Yao Yao and Dr. Zachary McEachin for providing sense and control ASOs, respectively.

## Author contributions

JP performed and analyzed all *in vitro* experiments. SA performed IHC in patient tissues. CZ, KT, DCP and GC helped in cell-based experiments and analysis. JP and JJ wrote the manuscript.

## Data and materials availability

All data are available in the main text or the supplementary materials.

## Supplementary figures

**Figure S1.**
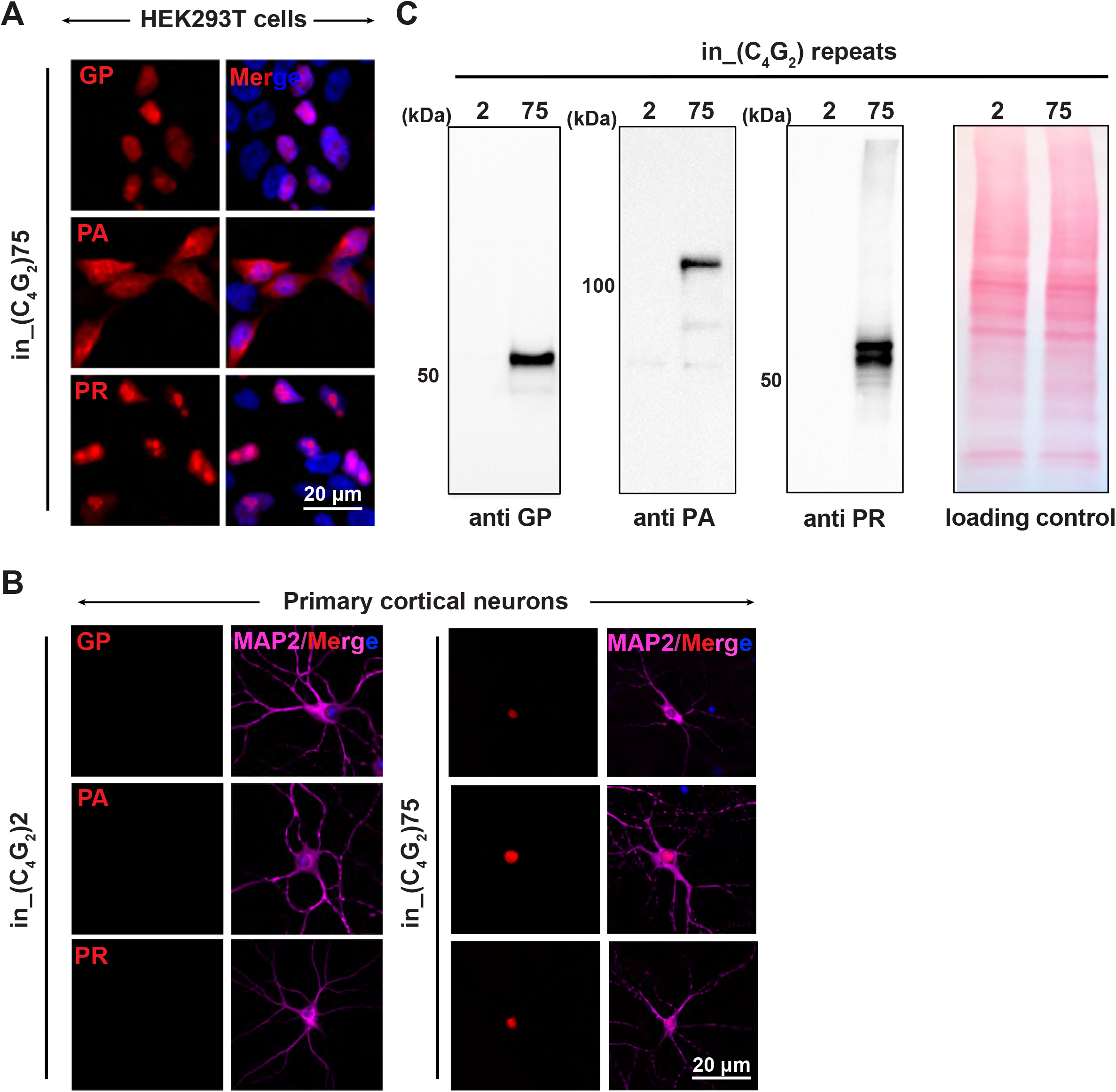
C9orf72 C_4_G_2_expanded repeats produce antisense DPR proteins in HEK293T cells and primary neurons. Representative images of DPR protein staining in (**A)** HEK293T and (**B)** primary neurons expressing in_(C_4_G_2_)75 repeats. Red, GP, PA, and PR; blue, DAPI; MAP2, Magenta. (**C)** Immunoblotting of DPR proteins in HEK293T expressing in_(C_4_G_2_)75 repeats. Ponceau staining was used as a loading control.

**Figure S2.**
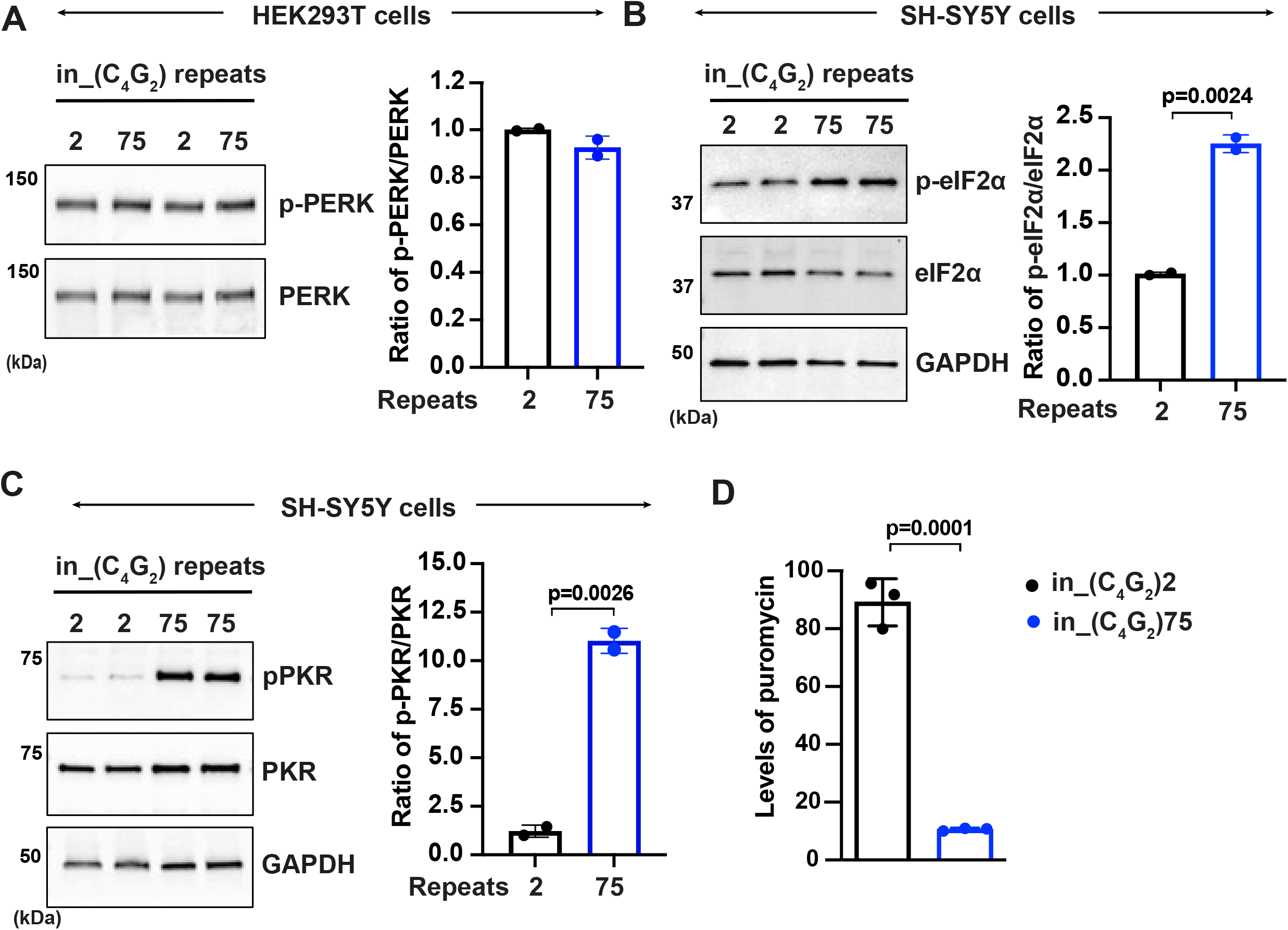
C9orf72 C_4_G_2_expanded repeats activate PKR/eIF2α-dependent integrated stress response in SH-SY5Y cells. **(A)** Immunoblotting of p-PERK in HEK293T cells expressing in_(C_4_G_2_)75 or (C_4_G_2_)2 repeats. Phosphorylated PERK levels were quantified and normalized to total PERK. GAPDH was used as a loading control. Error bars represent S.D. (n=2 independent experiments). Statistical analyses were performed using student’s t-test. **(B-C)** Immunoblotting of p-PKR and peIF2α in SH-SY5Y cells expressing (C_4_G_2_)75 or (C_4_G_2_)2 repeats. p-PKR (T446) and peIF2α (Ser51) were normalized to total PKR and eIF2α respectively. GAPDH was used as a loading control. Error bars represent S.D. (n=2 independent experiments). Statistical analyses were performed using student’s t-test. **(D)** Quantification of puromycin levels in HEK293T cells expressing in_(C_4_G_2_)75 or (C_4_G_2_)2 repeats. The level of puromycin was normalized to GAPDH. Error bars represent S.D. (n=3 independent experiments). Statistical analyses were performed using student’s t-test.

**Figure S3.**
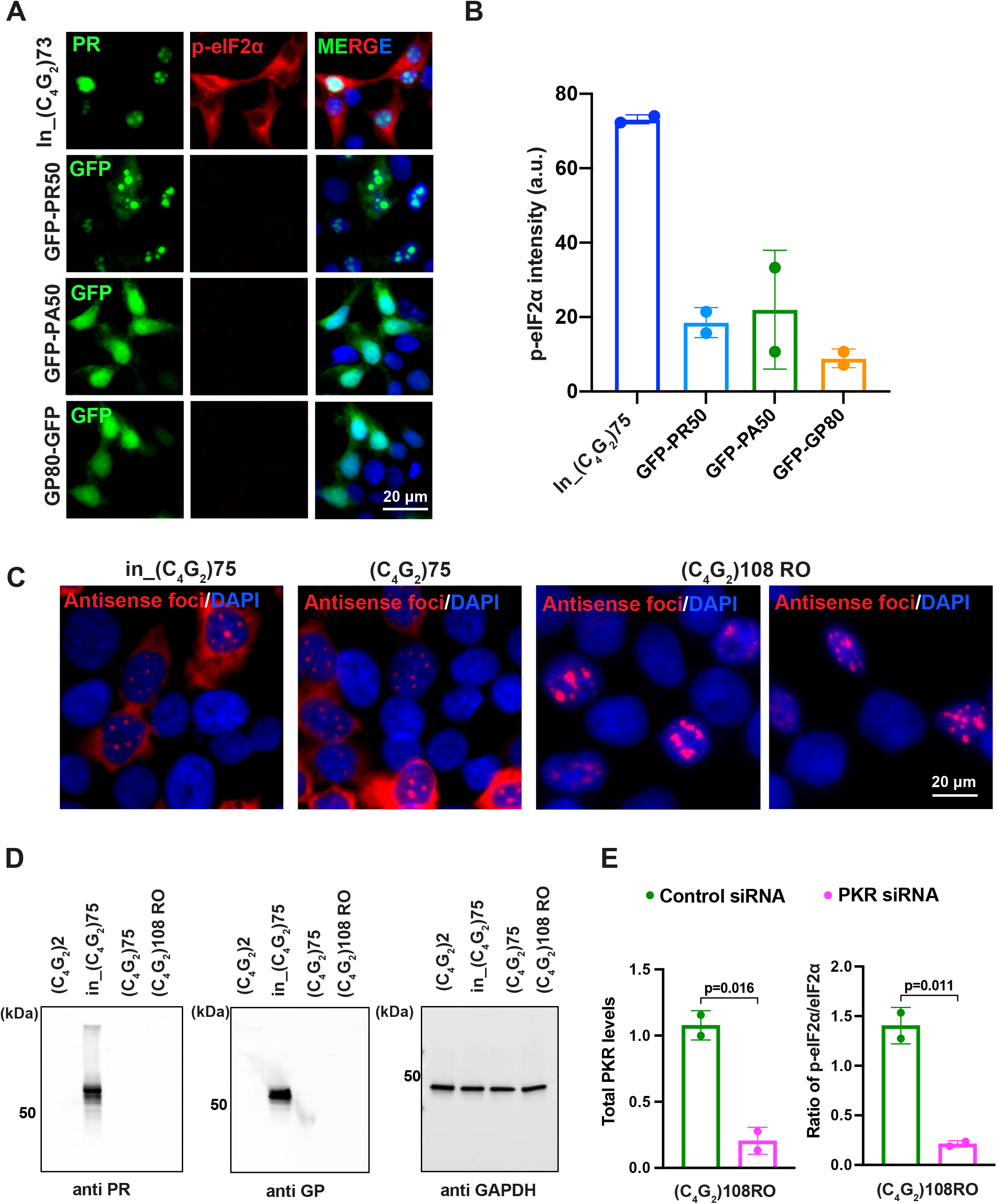
Antisense DPR proteins do not activate PKR/eIF2α-dependent integrated stress response. **(A-B)** Representative images **(A)** and quantification (**B)** of p-eIF2α staining in HEK293T cells expressing in_(C_4_G_2_)75, PR50, PA50, or GP80. Green, GP, PA, or PR; red, p-eIF2α; blue, DAPI. Error bars represent S.D. (n=2 independent experiments). **(C)** Representative images of antisense RNA foci in HEK293T cells expressing in_(C_4_G_2_)75, (C_4_G_2_)75 or (C_4_G_2_)108RO repeats. Foci were detected by RNA FISH. Red, foci; blue, DAPI. **(D)** Immunoblotting of DPR proteins in HEK293T cells expressing in_(C_4_G_2_)75, (C_4_G_2_)75 and (C_4_G_2_)108RO repeats. DPR protein levels were detected using anti-PR and anti-GP. GAPDH was used as a loading control. **(E)** Immunoblotting of PKR and p-eIF2α (Ser51) in HEK293T cells expressing (C_4_G_2_)108RO together with control or PKR siRNA. PKR and p-eIF2α (Ser51) were normalized to GADPH and total eIF2α respectively. Error bars represent S.D. (n=2 independent experiments). Statistical analyses were performed using student’s t-test.

**Figure S4.**
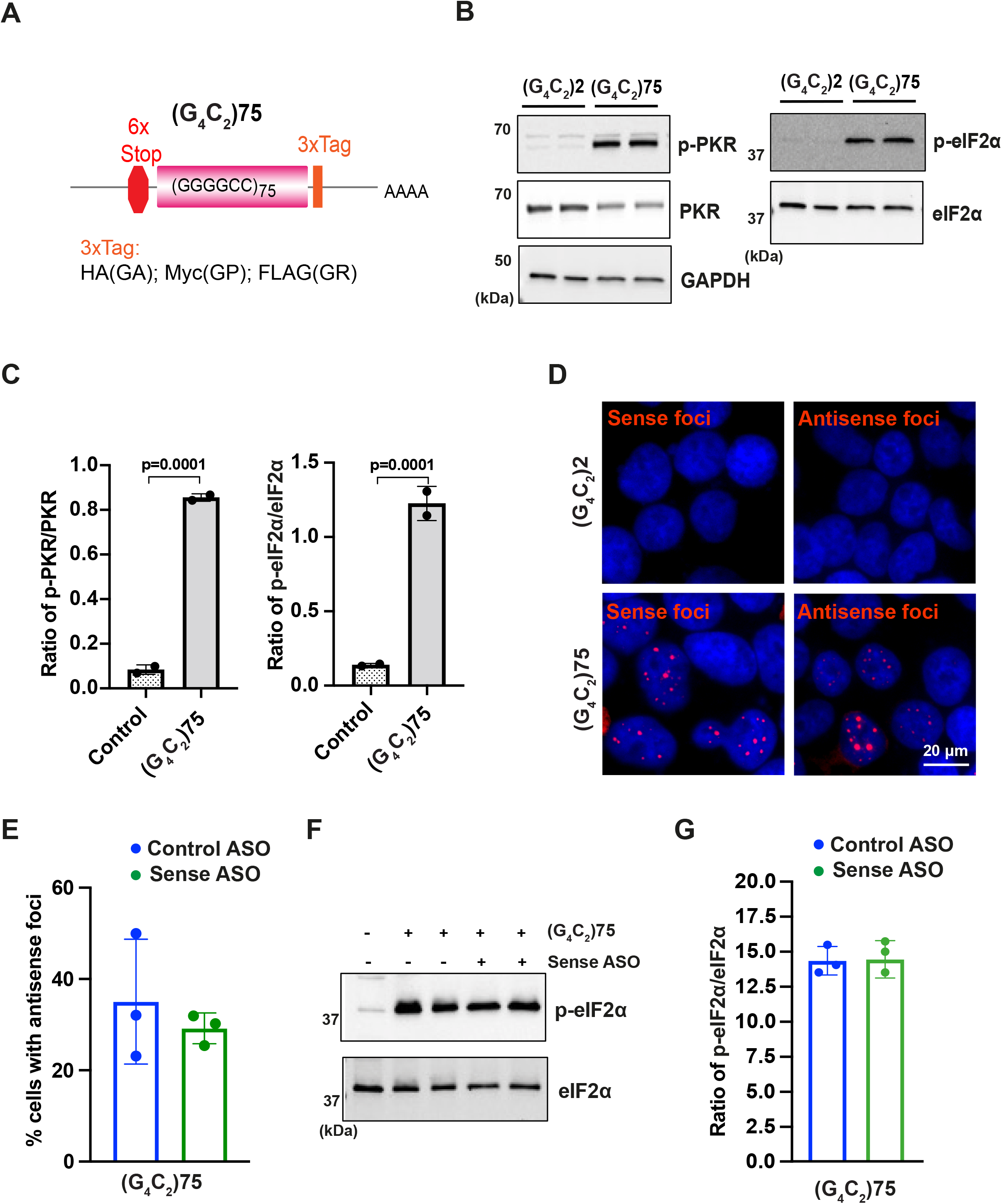
C9orf72 sense G_4_C_2_repeat expanded RNAs cannot activate the PKR/eIF2α pathway. (A) Schematic illustration of the (G_4_C_2_)75 repeat construct with 6× stop codons at the N-terminus 3× protein tags at the C-terminus of the repeats to monitor the DPR proteins in each frame. (**B-C)** Immunoblotting of p-PKR and p-eIF2α in HEK293T cells expressing (G_4_C_2_)75 or (G_4_C_2_)2 repeats. p-PKR (T446) and p-eIF2α (Ser51) were normalized to total PKR and eIF2α respectively. GAPDH was used as a loading control. Error bars represent S.D. (n=2 independent experiments). Statistical analyses were performed using student’s t-test. **(D)** Representative images of sense and antisense RNA foci in HEK293T cells expressing (G_4_C_2_)75. Foci were detected by RNA FISH. Red, foci; blue, DAPI. (**E)** Quantification of sense RNA foci in HEK293T expressing (G_4_C_2_)75 together with either control ASO or ASOs targeting sense (G_4_C_2_) repeat expanded RNAs. Error bars represent S.D. (n=80-100 cells/condition from 3 independent experiments). (**F)** Immunoblotting of p-eIF2α in HEK293T cells expressing (G_4_C_2_)75 or (G_4_C_2_)2 repeats together with either control ASO or ASOs targeting sense (G_4_C_2_) repeat expanded RNAs. **(G)** p-eIF2α (Ser51) was normalized to total eIF2α. GAPDH was used as a loading control. Error bars represent S.D. (n=3 independent experiments).

## References

1. Renton, A.E., et al., A hexanucleotide repeat expansion in C9ORF72 is the cause of chromosome 9p21-linked ALS-FTD. Neuron, 2011. 72(2): p. 257–68.

2. DeJesus-Hernandez, M., et al., Expanded GGGGCC hexanucleotide repeat in noncoding region of C9ORF72 causes chromosome 9p-linked FTD and ALS. Neuron, 2011. 72(2): p. 245–56.

3. La Spada, A.R. and J.P. Taylor, Repeat expansion disease: progress and puzzles in disease pathogenesis. Nat Rev Genet, 2010. 11(4): p. 247–58.

4. Jiang, J. and J. Ravits, Pathogenic Mechanisms and Therapy Development for C9orf72 Amyotrophic Lateral Sclerosis/Frontotemporal Dementia. Neurotherapeutics, 2019. 16(4): p. 1115–1132.

5. van Blitterswijk, M., et al., Novel clinical associations with specific C9ORF72 transcripts in patients with repeat expansions in C9ORF72. Acta Neuropathol, 2015. 130(6): p. 863–76.

6. Gijselinck, I., et al., A C9orf72 promoter repeat expansion in a Flanders-Belgian cohort with disorders of the frontotemporal lobar degeneration-amyotrophic lateral sclerosis spectrum: a gene identification study. Lancet Neurol, 2012. 11(1): p. 54–65.

7. Ash, P.E., et al., Unconventional translation of C9ORF72 GGGGCC expansion generates insoluble polypeptides specific to c9FTD/ALS. Neuron, 2013. 77(4): p. 639–46.

8. McEachin, Z.T., et al., Chimeric Peptide Species Contribute to Divergent Dipeptide Repeat Pathology in c9ALS/FTD and SCA36. Neuron, 2020. 107(2): p. 292-305.e6.

9. Tabet, R., et al., CUG initiation and frameshifting enable production of dipeptide repeat proteins from ALS/FTD C9ORF72 transcripts. Nat Commun, 2018. 9(1): p. 152.

10. Gao, F.B., J.D. Richter, and D.W. Cleveland, Rethinking Unconventional Translation in Neurodegeneration. Cell, 2017. 171(5): p. 994–1000.

11. Swinnen, B., et al., A zebrafish model for C9orf72 ALS reveals RNA toxicity as a pathogenic mechanism. Acta Neuropathol, 2018. 135(3): p. 427–443.

12. Therrien, M., et al., Deletion of C9ORF72 results in motor neuron degeneration and stress sensitivity in C. elegans. PLoS One, 2013. 8(12): p. e83450.

13. Koppers, M., et al., C9orf72 ablation in mice does not cause motor neuron degeneration or motor deficits. Ann Neurol, 2015. 78(3): p. 426–38.

14. Atanasio, A., et al., C9orf72 ablation causes immune dysregulation characterized by leukocyte expansion, autoantibody production, and glomerulonephropathy in mice. Sci Rep, 2016. 6: p. 23204.

15. Sudria-Lopez, E., et al., Full ablation of C9orf72 in mice causes immune system-related pathology and neoplastic events but no motor neuron defects. Acta Neuropathol, 2016. 132(1): p. 145–7.

16. O’Rourke, J.G., et al., C9orf72 is required for proper macrophage and microglial function in mice. Science, 2016. 351(6279): p. 1324–9.

17. Sullivan, P.M., et al., The ALS/FTLD associated protein C9orf72 associates with SMCR8 and WDR41 to regulate the autophagy-lysosome pathway. Acta Neuropathol Commun, 2016. 4(1): p. 51.

18. Ugolino, J., et al., Loss of C9orf72 Enhances Autophagic Activity via Deregulated mTOR and TFEB Signaling. PLoS Genet, 2016. 12(11): p. e1006443.

19. Burberry, A., et al., Loss-of-function mutations in the C9ORF72 mouse ortholog cause fatal autoimmune disease. Sci Transl Med, 2016. 8(347): p. 347ra93.

20. Harms, M.B., et al., Lack of C9ORF72 coding mutations supports a gain of function for repeat expansions in amyotrophic lateral sclerosis. Neurobiol Aging, 2013. 34(9): p. 2234 e13–9.

21. Chew, J., et al., Neurodegeneration. C9ORF72 repeat expansions in mice cause TDP-43 pathology, neuronal loss, and behavioral deficits. Science, 2015. 348(6239): p. 1151–4.

22. Jiang, J., et al., Gain of Toxicity from ALS/FTD-Linked Repeat Expansions in C9ORF72 Is Alleviated by Antisense Oligonucleotides Targeting GGGGCC-Containing RNAs. Neuron, 2016. 90(3): p. 535–50.

23. Choi, S.Y., et al., C9ORF72-ALS/FTD-associated poly(GR) binds Atp5a1 and compromises mitochondrial function in vivo. Nat Neurosci, 2019. 22(6): p. 851–862.

24. Mizielinska, S., et al., C9orf72 repeat expansions cause neurodegeneration in Drosophila through arginine-rich proteins. Science, 2014. 345(6201): p. 1192–1194.

25. Zhang, Y.J., et al., Heterochromatin anomalies and double-stranded RNA accumulation underlie C9orf72 poly(PR) toxicity. Science, 2019. 363(6428).

26. Zhu, Q., et al., Reduced C9ORF72 function exacerbates gain of toxicity from ALS/FTD-causing repeat expansion in C9orf72. Nat Neurosci, 2020. 23(5): p. 615–624.

27. Boivin, M., et al., Reduced autophagy upon C9ORF72 loss synergizes with dipeptide repeat protein toxicity in G4C2 repeat expansion disorders. EMBO J, 2020. 39(4): p. e100574.

28. Donnelly, C.J., et al., RNA toxicity from the ALS/FTD C9ORF72 expansion is mitigated by antisense intervention. Neuron, 2013. 80(2): p. 415–28.

29. Sareen, D., et al., Targeting RNA foci in iPSC-derived motor neurons from ALS patients with a C9ORF72 repeat expansion. Sci Transl Med, 2013. 5(208): p. 208ra149.

30. Lee, Y.B., et al., Hexanucleotide repeats in ALS/FTD form length-dependent RNA foci, sequester RNA binding proteins, and are neurotoxic. Cell Rep, 2013. 5(5): p. 1178–86.

31. Mori, K., et al., hnRNP A3 binds to GGGGCC repeats and is a constituent of p62-positive/TDP43-negative inclusions in the hippocampus of patients with C9orf72 mutations. Acta Neuropathol, 2013. 125(3): p. 413–23.

32. Xu, Z., et al., Expanded GGGGCC repeat RNA associated with amyotrophic lateral sclerosis and frontotemporal dementia causes neurodegeneration. Proc Natl Acad Sci U S A, 2013. 110(19): p. 7778–83.

33. Haeusler, A.R., et al., C9orf72 nucleotide repeat structures initiate molecular cascades of disease. Nature, 2014. 507(7491): p. 195–200.

34. Conlon, E.G., et al., The C9ORF72 GGGGCC expansion forms RNA G-quadruplex inclusions and sequesters hnRNP H to disrupt splicing in ALS brains. Elife, 2016. 5.

35. Celona, B., et al., Suppression of C9orf72 RNA repeat-induced neurotoxicity by the ALS-associated RNA-binding protein Zfp106. Elife, 2017. 6.

36. Wen, X., et al., Antisense proline-arginine RAN dipeptides linked to C9ORF72-ALS/FTD form toxic nuclear aggregates that initiate in vitro and in vivo neuronal death. Neuron, 2014. 84(6): p. 1213–25.

37. Tao, Z., et al., Nucleolar stress and impaired stress granule formation contribute to C9orf72 RAN translation-induced cytotoxicity. Hum Mol Genet, 2015. 24(9): p. 2426–41.

38. Lee, K.H., et al., C9orf72 Dipeptide Repeats Impair the Assembly, Dynamics, and Function of Membrane-Less Organelles. Cell, 2016. 167(3): p. 774–788 e17.

39. Zu, T., et al., Non-ATG-initiated translation directed by microsatellite expansions. Proc Natl Acad Sci U S A, 2011. 108(1): p. 260–5.

40. Kanekura, K., et al., Poly-dipeptides encoded by the C9ORF72 repeats block global protein translation. Hum Mol Genet, 2016. 25(9): p. 1803–13.

41. Zhang, Y.J., et al., Aggregation-prone c9FTD/ALS poly(GA) RAN-translated proteins cause neurotoxicity by inducing ER stress. Acta Neuropathol, 2014. 128(4): p. 505–24.

42. May, S., et al., C9orf72 FTLD/ALS-associated Gly-Ala dipeptide repeat proteins cause neuronal toxicity and Unc119 sequestration. Acta Neuropathol, 2014. 128(4): p. 485–503.

43. Yamakawa, M., et al., Characterization of the dipeptide repeat protein in the molecular pathogenesis of c9FTD/ALS. Hum Mol Genet, 2015. 24(6): p. 1630–45.

44. Freibaum, B.D., et al., GGGGCC repeat expansion in C9orf72 compromises nucleocytoplasmic transport. Nature, 2015. 525(7567): p. 129–33.

45. Yang, D., et al., FTD/ALS-associated poly(GR) protein impairs the Notch pathway and is recruited by poly(GA) into cytoplasmic inclusions. Acta Neuropathol, 2015. 130(4): p. 525–35.

46. Boeynaems, S., et al., Drosophila screen connects nuclear transport genes to DPR pathology in c9ALS/FTD. Sci Rep, 2016. 6: p. 20877.

47. Zhang, Y.J., et al., C9ORF72 poly(GA) aggregates sequester and impair HR23 and nucleocytoplasmic transport proteins. Nat Neurosci, 2016. 19(5): p. 668–677.

48. Schludi, M.H., et al., Spinal poly-GA inclusions in a C9orf72 mouse model trigger motor deficits and inflammation without neuron loss. Acta Neuropathol, 2017. 134(2): p. 241–254.

49. Zhang, Y.J., et al., Poly(GR) impairs protein translation and stress granule dynamics in C9orf72-associated frontotemporal dementia and amyotrophic lateral sclerosis. Nat Med, 2018. 24(8): p. 1136–1142.

50. Hao, Z., et al., Motor dysfunction and neurodegeneration in a C9orf72 mouse line expressing poly-PR. Nat Commun, 2019. 10(1): p. 2906.

51. Moens, T.G., et al., Sense and antisense RNA are not toxic in Drosophila models of C9orf72-associated ALS/FTD. Acta Neuropathol, 2018. 135(3): p. 445–457.

52. Tran, H., et al., Differential Toxicity of Nuclear RNA Foci versus Dipeptide Repeat Proteins in a Drosophila Model of C9ORF72 FTD/ALS. Neuron, 2015. 87(6): p. 1207–1214.

53. Zhang, K., et al., The C9orf72 repeat expansion disrupts nucleocytoplasmic transport. Nature, 2015. 525(7567): p. 56–61.

54. Mizielinska, S., et al., C9orf72 frontotemporal lobar degeneration is characterised by frequent neuronal sense and antisense RNA foci. Acta Neuropathol, 2013. 126(6): p. 845–57.

55. DeJesus-Hernandez, M., et al., In-depth clinico-pathological examination of RNA foci in a large cohort of C9ORF72 expansion carriers. Acta Neuropathol, 2017. 134(2): p. 255–269.

56. Vatsavayai, S.C., et al., C9orf72-FTD/ALS pathogenesis: evidence from human neuropathological studies. Acta Neuropathol, 2019. 137(1): p. 1–26.

57. McEachin, Z.T., et al., RNA-mediated toxicity in C9orf72 ALS and FTD. Neurobiol Dis, 2020. 145: p. 105055.

58. Sellier, C., et al., Translation of Expanded CGG Repeats into FMRpolyG Is Pathogenic and May Contribute to Fragile X Tremor Ataxia Syndrome. Neuron, 2017. 93(2): p. 331–347.

59. He, F., et al., The carboxyl termini of RAN translated GGGGCC nucleotide repeat expansions modulate toxicity in models of ALS/FTD. Acta Neuropathol Commun, 2020. 8(1): p. 122.

60. Zu, T., et al., RAN proteins and RNA foci from antisense transcripts in C9ORF72 ALS and frontotemporal dementia. Proc Natl Acad Sci U S A, 2013. 110(51): p. E4968–77.

61. Handa, V., T. Saha, and K. Usdin, The fragile X syndrome repeats form RNA hairpins that do not activate the interferon-inducible protein kinase, PKR, but are cut by Dicer. Nucleic Acids Res, 2003. 31(21): p. 6243–8.

62. Tian, B., et al., Expanded CUG repeat RNAs form hairpins that activate the double-stranded RNA-dependent protein kinase PKR. Rna, 2000. 6(1): p. 79–87.

63. Martinez, N.W., F.E. Gomez, and S. Matus, The Potential Role of Protein Kinase R as a Regulator of Age-Related Neurodegeneration. Front Aging Neurosci, 2021. 13: p. 638208.

64. Schmidt, E.K., et al., SUnSET, a nonradioactive method to monitor protein synthesis. Nat Methods, 2009. 6(4): p. 275–7.

65. Green, K.M., et al., RAN translation at C9orf72-associated repeat expansions is selectively enhanced by the integrated stress response. Nat Commun, 2017. 8(1): p. 2005.

66. Zu, T., et al., Metformin inhibits RAN translation through PKR pathway and mitigates disease in <em>C9orf72</em> ALS/FTD mice. Proceedings of the National Academy of Sciences, 2020: p. 202005748.

67. Rodriguez, S., et al., Genome-encoded cytoplasmic double-stranded RNAs, found in C9ORF72 ALS-FTD brain, propagate neuronal loss. Sci Transl Med, 2021. 13(601).

68. Zu, T., et al., Metformin inhibits RAN translation through PKR pathway and mitigates disease in C9orf72 ALS/FTD mice. Proc Natl Acad Sci U S A, 2020. 117(31): p. 18591–18599.

69. Jovicic, A., et al., Modifiers of C9orf72 dipeptide repeat toxicity connect nucleocytoplasmic transport defects to FTD/ALS. Nat Neurosci, 2015. 18(9): p. 1226–9.

70. Aladesuyi Arogundade, O., et al., Antisense RNA foci are associated with nucleoli and TDP-43 mislocalization in C9orf72-ALS/FTD: a quantitative study. Acta Neuropathol, 2019. 137(3): p. 527–530.

71. Cooper-Knock, J., et al., Antisense RNA foci in the motor neurons of C9ORF72-ALS patients are associated with TDP-43 proteinopathy. Acta Neuropathol, 2015. 130(1): p. 63–75.

72. Kovanda, A., et al., Anti-sense DNA d(GGCCCC)n expansions in C9ORF72 form i-motifs and protonated hairpins. Sci Rep, 2015. 5: p. 17944.

73. Reddy, K., et al., The disease-associated r(GGGGCC)n repeat from the C9orf72 gene forms tract length-dependent uni-and multimolecular RNA G-quadruplex structures. J Biol Chem, 2013. 288(14): p. 9860–6.

74. Fratta, P., et al., C9orf72 hexanucleotide repeat associated with amyotrophic lateral sclerosis and frontotemporal dementia forms RNA G-quadruplexes. Sci Rep, 2012. 2: p. 1016.

75. Sadeq, S., et al., Endogenous Double-Stranded RNA. Noncoding RNA, 2021. 7(1).

76. Prudencio, M., et al., Repetitive element transcripts are elevated in the brain of C9orf72 ALS/FTLD patients. Hum Mol Genet, 2017. 26(17): p. 3421–3431.

77. Cheng, W., et al., C9ORF72 GGGGCC repeat-associated non-AUG translation is upregulated by stress through eIF2alpha phosphorylation. Nat Commun, 2018. 9(1): p. 51.

78. Valdez-Sinon, A.N., et al., Cdh1-APC Regulates Protein Synthesis and Stress Granules in Neurons through an FMRP-Dependent Mechanism. iScience, 2020. 23(5): p. 101132.

79. Weskamp, K., et al., Monitoring Neuronal Survival via Longitudinal Fluorescence Microscopy. J Vis Exp, 2019(143).

